# Mutant p53^R175H^-Associated Protection of Ovarian Cancer Precursor Cells from Cisplatin Requires Expression of SLC6A6 Taurine Transporter

**DOI:** 10.1101/2025.01.03.631237

**Authors:** Teagan Polotaye, Daniel Centeno, Marcin Iwanicki

## Abstract

Tumor suppressor gene TP53 is ubiquitously mutated in ovarian cancer precursor lesions that undergo pervasive accumulation of DNA damage. Prior literature provides evidence that the expression of mutant p53 protein in epithelial cells is associated with increased survival in response to DNA-damaging treatments. Hence, identification and understanding of the mechanisms that, in response to DNA damage accumulation, support survival of ovarian cancer precursor cells carrying p53 mutations might provide important information about the evolution of the disease. Here we used a combination of OC precursor cell models, biochemistry, microscopy, and flow cytometry to provide evidence that the taurine transporter, the SLC6A6 molecule, contributes to cell protection from DNA-damaging (cisplatin) treatment. We found that expression of mutant p53^R175H^ in OC precursor cells, the fallopian tube non-ciliated epithelial (FNE) cells, induced resistance to the DNA-damaging agent cisplatin. Most importantly, shRNA-mediated targeting of SLC6A6 transcript re-sensitized FNE cells expressing mutant p53^R175H^ to cisplatin treatment. Our studies are consistent with the model that the loss of SLC6A6 alters mechanisms involved in the regulation of cell survival in response to DNA damage.

## INTRODUCTION

Most ovarian cancers (OCs) arise from the fallopian tube non-ciliated epithelial (FNE) cells that accumulate p53 mutations and DNA damage [1]. P53 protein is a major tumor suppressor molecule that regulates a plethora of cell functions including cell cycle progression, DNA damage response, metabolism, motility, and cell death [2] . In mouse models, loss of p53 function inevitably leads to genome instability, accumulation of DNA damage, loss of cell-cycle control, and cancer development [3] . In human OC, high-grade serous ovarian carcinoma, p53 mutations are nearly ubiquitous [4] implicating a cancer driver molecular event. Therefore, studying the role of p53 mutations in the context of OC precursor cells would provide an opportunity to understand the molecular mechanism of the disease development. In this manuscript we describe experiments to study the role of mutant p53 expression in OC precursor cells, the fallopian tube non-ciliated epithelial (FNE) cells.

While most of the OC cells respond well to initial rounds of DNA-damaging treatments, some OC cells can resist the therapy, and drive disease recurrence [5]. This suggests the possibility that selected OC precursor cells might possess intrinsic mechanisms that support survival under DNA-damaging conditions. In this manuscript, we demonstrate that the expression of mutant p53^R175H^ in FNE cells induces cell protection from cisplatin, the commonly used cancer therapeutic.

Taurine is a non-proteogenic amino acid implicated in cell protection [6]. Recent reports [7] including ours [8], demonstrated taurine’s ability to protect cells from DNA-damaging chemotherapeutic treatment. Furthermore, evidence exists that taurine transporter solute carrier family member 6 A6 (SLC6A6) is enriched in chemotherapy-resistant colorectal cancer (CRC) cell populations [9], and shRNA-mediated attenuation of SLC6A6 sensitizes a CRC cell line to chemotherapy [10]. This information motivated us to ask an important question whether targeting SLC6A6 in FNE cells expressing mutant p53^R175H^ promotes cisplatin sensitization. Our investigations revealed that mutant p53 protected FNE cells from cisplatin, and SLC6A6 significantly contributed to this resistance mechanism.

## RESULTS

### Phenotypic characterization of FNE cells depleted of SLC6A6

To assess the role of SLC6A6 in FNE cells expressing mutant p53 we used lentiviral delivery of small hairpin (sh) RNA containing SLC6A6 antisense. We first confirmed mutant p53 and SLC6A6 levels in FNE cells expressing wild-type (WT) or mutant p53 (**Fig.1A**). We then proceeded to SLC6A6 attenuation using previously published SLC6A6shRNA_G8 [8]. SLC6A6 is a membrane protein, thus we utilized a membrane protein extraction kit to measure SLC6A6 protein levels in various FNE cells expressing control (scramble) or SLC6A6 targeting hairpin. We found, using western blot, that lentiviral transduction of SLC6A6 shRNA but not scramble shRNA resulted in SLC6A6 protein expression attenuation in control (WT p53) FNE cells (**Fig. 1B**), and FNE cells expressing mutant p53^R175H^ (**Fig. 1C**).

**Figure 1.**
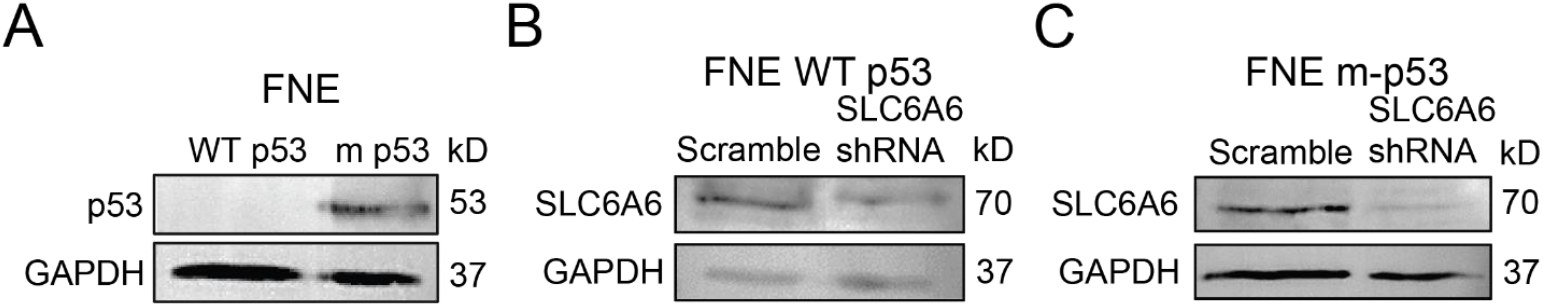
(A) p53 levels in FNE cells expressing WT p53 or m p53. (B) SLC6A6 levels in FNE WT p53 cells treated with scramble or shRNA vector. (C) SLC6A6 levels in FNE WT p53 cells treated with scramble or shRNA vector.

Next, we used phase contrast microscopy recording and cell counting to determine whether SLC6A6 depletion resulted in changes in cell morphology, and/or proliferation. We found that SLC6A6 knockdown increased cell area in FNE cells expressing WT but not mutant p53^R175H^ (**Fig. 2A-B**), without affecting proliferation (**Fig. 2C**) indicating a possibility that loss of SLC6A6 activates WT p53-dependent mechanisms related to regulation of cell morphology. While depletion of SLC6A6 from FNE cells expressing mutant p53^R175H^ had no effect on cell morphology (**Fig. 2D-E**), it slightly induced proliferation (**Fig. 2F**).

**Figure 2.**
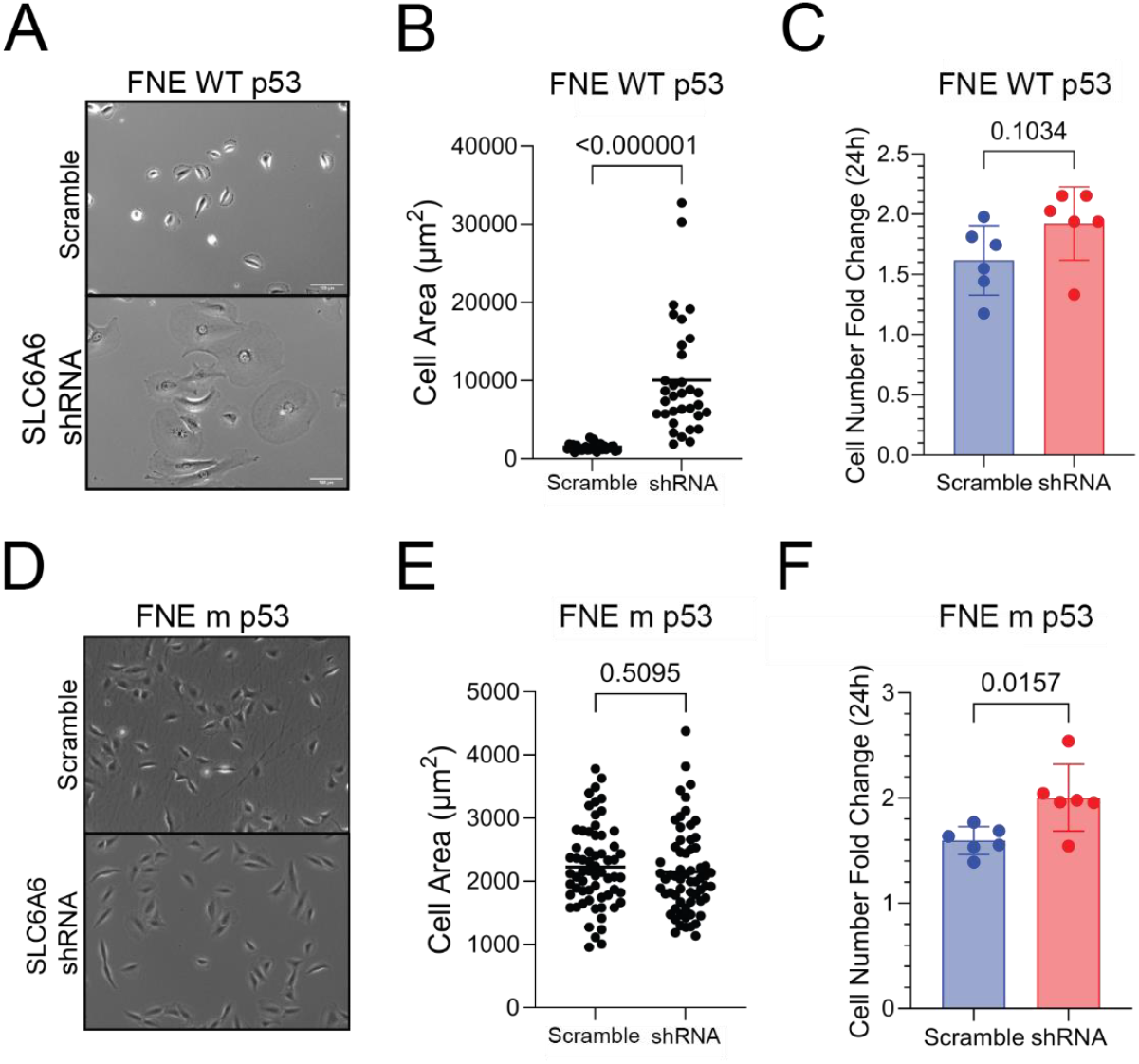
(A) Phase contrast image of FNE WT p53 cells expressing scramble or SLC6A6 shRNA vector. Scale bar is 100 μm. (B) Quantification of FNE WT p53 cell area. Data is reported as mean ± SD and each dot represents one cell. Significance tested using unpaired t-test. (C) 24-hour cell number fold change from phase contrast images. Data is normalized to 0-hour time point and is reported as mean ± SD. Each dot represents one replicate. Significance tested using t-test. (D) Phase contrast image of FNE m p53 cells expressing scramble or SLC6A6 shRNA vector. Scale bar is 100 μm. (E) Quantification of FNE m p53 cell area. Data is reported as mean ± SD and each dot represents one cell. Significance tested using unpaired t-test. (F) 24-hour cell number fold change from phase contrast images. Data is normalized to 0-hour time point and is reported as mean ± SD. Each dot represents one replicate. Significance tested using t-test.

### Depletion of SLC6A6 sensitizes FNE cells expressing mutant p53^R175H^ to cisplatin

Up to this point, our data provide evidence that under normal culture conditions, the ability of SLC6A6 to regulate cell morphology and cell proliferation is linked to p53 status. Next, we wanted to explore whether p53 status (WT vs mutant p53) also determines SLC6A6 contribution to cell survival under DNA-damaging stress. Thus, we exposed FNE cells (WT and mutant p53) expressing control or SLC6A6 shRNA to 10 μM cisplatin for 72 hours. Following cisplatin exposure, we used a combination of flow cytometry and cell death detection propidium iodide-based statins to determine cell viability. Our study revealed the following results. First, depletion of SLC6A6 did not cause cell death in FNE cells expressing WT or mutant p53 (**Fig. 3A-B**). Second, consistent with the literature [11] mutant p53 significantly decreased FNE cell cisplatin sensitivity (**Fig. 3A-B)**. Third, depletion of SLC6A6 increased the cisplatin sensitivity of FNE cells expressing WT or mutant p53 (**Fig. 3 A-B)**. This experiment provided evidence that SLC6A6 can mediate cisplatin resistance in OC precursor cells carrying cancer-driving mutations.

**Figure 3.**
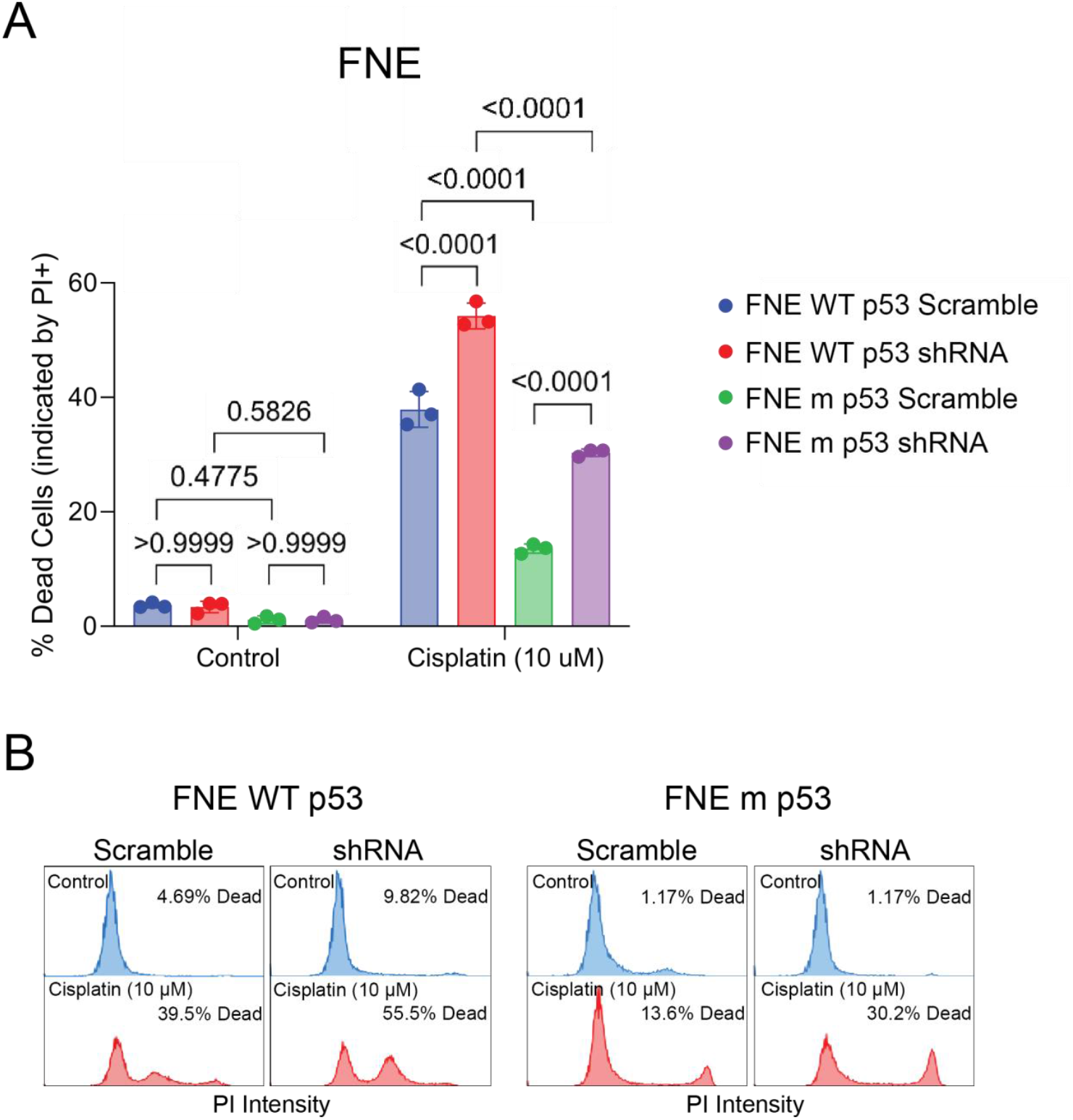
(A) Flow cytometric analysis of PI+ FNE cells expressing WT or m p53 in response to 10μM cisplatin. Data is reported as mean ± SD and each dot represents one replicate. Significance tested using 2-way ANOVA. (B) Histogram showing concatenated replicates from (A) and average PI+ signal.

## DISCUSSION

In this study, we used a combination of FNE cell cultures, lentiviral-mediated manipulation of gene expression, microscopy, western blotting, and flow cytometry to provide important information about the role of SLC6A6 in the regulation of cell proliferation, cell size, and response to cisplatin. We report that the depletion of SLC6A6 induces cell size increase, regulates cell proliferation, and promotes cisplatin sensitivity. While the effect of SLC6A6 depletion on morphology, and cell proliferation was linked to p53 status, the cisplatin sensitization was not, indicating that SLC6A6 can regulate cell survival of FNE cells expressing cancer-driving p53 mutations. How SLC6A6 modulates cell size and cell proliferation is not known. However, our data suggest a link between SLC6A6 and WT p53 expression. SLC6A6 has been reported to localize to the nucleus[12], raising the possibility of regulating molecular mechanisms that take place in the nucleus, for instance, modulation of p53 transcriptional activity. Furthermore, it has previously been reported that the depletion of SLC6A6 induces cell-cycle arrest [6], a phenotype associated with cell size increase, and irreversible cell cycle arrest. However, data presented in this report, indicate that, in FNE cells expressing WT p53, SLC6A6 impacts morphology but not cell proliferation. So, we hypothesize that the loss of SLC6A6 activates WT p53 to induce changes in cell size but not cell proliferation. In follow-up experiments, we will examine this hypothesis by questioning whether depletion of SLC6A6 activates p53 target genes, implicate in cell size regulation, in FNE cells expressing WT p53.

SLC6A6 attenuation in FNE cells expressing WT or mutant p53 led to cisplatin sensitization, indicating that SLC6A6 might modulate DNA damage response independently of p53 status. Recent studies demonstrated that genetic ablation of SLAC6A6 can lead to increased DNA damage, and sensitivity to DNA-damaging compounds. These studies raised the question about possible SLC6A6 involvement in the DNA repair process. Whether and how SLC6A6 impacts DNA repair is not currently known. However, evidence suggests that the amino acid taurine, which is transported into cells by SLC6A6, decreases DNA damage [6], and activates p53 and DNA-damage sensing mechanisms including protein kinases DNA-PK and ATM/ATR [8]. Thus, it is possible that SLC6A6-mediated taurine transport can modulate DNA repair.

In conclusion, our experiments shed new light on the taurine transporter, SLC6A6, as a contributor to cellular resistance mechanisms to the chemotherapy agent cisplatin. We have demonstrated that SLC6A6 has the potential to serve as a therapeutic target to increase cisplatin sensitivity.

## MATERIALS & METHODS

### Cell culture

FNE WT p53 and FNE m p53 cells were cultured in a 1:1 ratio of Medium 199 (HiMedia) and DMEM/F12 (HiMedia) supplemented with 2% inactivated Fetal Bovine serum (HI-FBS), 1% v/v penicillin-streptomycin (VWR), 0.5 ng/mL of beta-estradiol (US Biological), 14 μg/mL of insulin (Sigma-Aldrich), 0.2 pg/mL of triiodothyronine (Sigma-Aldrich), 0.025 μg/mL of all-trans retinoic acid (Beantown Chemical), 0.5 μg hydrocortisone (Sigma-Aldrich), 0.5 ng/mL of EGF (Peprotech), and 25 ng/mL of cholera toxin (Calbiochem). The cells were cultured within a humidified incubator at 37°C and with 5% carbon dioxide.

### Plasmids

To knockdown SLC6A6, a shRNA vector from Dharmacon (TRCN0000038412: TATACTTGTACTTGTTGTAGC) referred to as “SLC6A6shRNA_G8” in the manuscript was inserted into a p.LKO backbone.

### Lentivirus production

To produce lentivirus, packaging plasmids psPAX2 and pMD2.G were mixed with a lentiviral plasmid containing the shRNA and incubated with Lipofectamine 3000 (Invitrogen) in serum-free Opti-MEM (Gibco) to generate liposomes containing the plasmid DNA. The plasmids were then added to HEK293T cells to generate the virus. Media supernatants containing viral particles were collected 48 hours and 72 hours after transfection. FNE cells were then transduced by adding viral particles to cell culture media containing polybrene (Santa Cruz Biotechnology) at a concentration of 10 μg/mL and selected with puromycin 24 hours after transduction to generate cells that stably express the shRNA vector.

### Western blot

Cells were seeded and cultured in 10-cm dishes until ∼75% confluent, then treated with indicated treatment for the indicated time. The cells were then harvested via trypsinization and spun down at 300 x g for 5 minutes. Using the materials and protocol provided by the Mem-Per Plus Membrane Protein Extraction Kit (Thermo Fisher) supplemented with Halt TM Protease Inhibitor Cocktail (Thermo Fisher), cytosolic and membrane protein lysates were collected. The lysates were quantified via BCA Assay (Pierce) according to the provided protocol. The lysates were then mixed with a 6X sample buffer (375 mM Tris Base, 9% sodium dodecyl sulfate, 50% glycerol, 0.075% bromophenol blue, and 9% β-mercaptoethanol). The cytosolic lysates were boiled at 95°C for 10 minutes, and the membrane lysates were incubated in a water bath at 37°C for 30 minutes. The indicated samples were then loaded on a 10% polyacrylamide gel and resolved via electrophoresis. Proteins were then transferred from the gel to an Immobilon PVDF membrane (Immobilion), blocked with 5% w/v non-fat milk in Tris-buffered saline containing 0.1% v/v Tween-20 (TBST) for an hour at room temperature with constant mixing. Membranes were then incubated overnight at 4°C in primary antibodies diluted in 5% non-fat milk in TBST with constant mixing. The following day the membranes were washed three times with TBST and incubated in HRP-conjugated secondary antibody (1:10,000) for an hour at room temperature with constant mixing. The membranes were once again washed three times with TBST for ten minutes each. Membranes were then developed using Immoblion TM Forte enhanced chemiluminescent substrate (Millipore) and visualized using an iBright CL1500 (Thermo Fisher). For SLC6A6, Anti-SLC6A6/TauT (Abcam ab236898) was used at a dilution factor of 1:1000. For GAPDH, GAPDH (0411) (Santa Cruz sc-47724) was used at a dilution factor of 1:1000. For p53, p53 (Cell Signaling 2527S) was used at a dilution factor of 1:2000.

### Cell area measurement

Images were taken using an Agilent BioTek Lionheart FX automated fluorescence microscope (BTLFXm) using the 10X objective in the phase contrast channel at six different regions of interest (ROI). Using the polygon tool in FIJI (FIJI is Just ImageJ), cell area was measured and converted from pixels^2^ to um^2^ using the provided measurements in the image metadata. Significance was tested using an unpaired t-test.

### Cell proliferation measurement

To measure cell proliferation, images were taken using the GFP channel with the 10X objective of an Agilent BioTek Lionheart FX automated fluorescence microscope at six different ROI at 0-hour and 24-hour time points. Images were analyzed using FIJI.

### Flow cytometry

To determine cell viability, cell culture media was collected, and adherent cells were harvested via trypsinization. The cells were centrifuged at 300 x g for 5 minutes, and then washed twice with PBS supplemented with 2% FBS. The cells were then incubated on ice in PBS supplemented with 2% FBS and 2 μg/mL propidium iodide. Data was acquired via flow cytometry using an Attune NxT Flow Cytometer and then analyzed using FlowJo software.

## Notes

### Competing Interest Statement

The authors have declared no competing interest.

### Summary of Updates

Updated formatting of PDF to stop image files from displaying on separate page.

